# Colony maintenance and the behavioral and physiological characteristics of selectively bred obesity prone and obesity resistant rats

**DOI:** 10.64898/2026.07.03.736414

**Authors:** Joy Sales Colquitt, Lauren M. Raycraft, Rowan J. Calkins, Maria Ortego-Dominguez, Carrie R. Ferrario

**Affiliations:** Department of Pharmacology, The University of Michigan, Ann Arbor, MI

**Keywords:** obesity models, genetic predisposition, obesity susceptibility, striatum, reward, motivation, nucleus accumbens, instrumental responding

## Abstract

Obesity arises from interactions between several factors including physiology, environment and genes. Studies in humans have revealed that up to 70% of overweight and obesity can be attributed to biological and genetic factors. Thus, rodent models that capture innate susceptibility or resistance to obesity have been invaluable for disentangling inherent drivers of obesity from neurobiological alterations that occur in response to consumption of obesogenic foods and/or increased adiposity. For example, studies of rats selectively bred for their propensity vs resistance to diet-induced weight gain (DIO and DR) have uncovered differences in hypothalamic circuits involved in leptin signaling and revealed relationships between susceptibility to obesity and motivational response to food cues, as well as inherent and diet-induced alterations in mesocorticolimbic systems that differ between these populations. Maintaining selectively bred lines in a closed breeding population requires the periodic introduction of new genes to avoid inbreeding. Here we describe a process for maintaining these lines, characterize key phenotypes across the selection process and verify weight gain and obesity phenotypes in the resulting colony. In addition, given the central role of the striatum in motivation for food, we examined basal striatal function and food motivation in these refreshed lines using whole-cell patch clamping and instrumental procedures. Key weight and metabolic phenotypes were maintained in the resulting colony, as was enhanced motivation for food in obesity prone rats. This provides a strong basis for examination of interactions between genes, environment and neurobehavioral plasticity that promote weight gain and obesity.

## Introduction

Obesity is a complex disease that arises from interactions between genetic, physiological, and environmental factors. In the United States, it is estimated that 2 in 5 adults and 1 in 5 children live with obesity and, as of 2022, overweight and obesity affected more than 2 billion people worldwide (Ward et al., 2019; Collaboration, 2024; Collaborators, 2024). Largescale studies in humans have revealed that up to 70% of overweight and obesity (based on BMI) can be attributed to biological and genetic factors that influence basal metabolic rate and neurophysiological systems that regulate hunger, satiety, and motivation for food (Maes et al., 1997; Fall and Ingelsson, 2014; Locke et al., 2015; Sandholt et al., 2015; van der Klaauw and Farooqi, 2015). Thus, rodent models have been developed that capture intrinsic susceptibility or resistance to obesity (Madsen et al., 2010; Lutz, 2018; Lommi et al., 2025). These models have been invaluable for disentangling inherent drivers of obesity from neurobiological alterations that occur in response to consumption of obesogenic foods and/or increased adiposity (Levin et al., 1997; Madsen et al., 2010; Patil and McCarroll, 2013; Lommi et al., 2025).

One such model was developed by Barry Levin and colleagues (1997) when they observed that there is naturally occurring variance in the propensity to gain weight in outbred Sprague Dawley rats when given *ad libitum* access to a moderately fatty, calorically dense diet. The Levin group selectively bred for the trait of high or low weight gain and established lines of rats that they referred to as diet-induced obese (DIO) and diet-resistant (DR, Levin et al., 1997). When maintained on standard lab chow, DIO rats tend to eat more than DR rats, and when given long-term free access to calorically dense diets, DIO rats develop pronounced obesity and metabolic disruption (e.g., Levin and Keesey, 1998; Vollbrecht et al., 2015; Alonso-Caraballo et al., 2018). In contrast, DR rats tend to be smaller and leaner when maintained on chow and have only modest increases in adiposity when maintained on calorically dense diets. Weight gain and increased adiposity in DIO rats is more pronounced in males than females, consistent with established sex differences in the regulation of appetite and weight gain (Hirschberg, 2012; Alonso-Caraballo et al., 2018).

Using these selectively bred rats, Levin and colleagues uncovered differences in hypothalamic circuits involved in leptin signaling between DIOs and DRs (Bouret et al., 2008; Scarpace and Zhang, 2009; Madsen et al., 2010) as well as interactions between maternal obesity and susceptibility to weight gain in adulthood (Levin, 2010b, a; Patterson et al., 2010). More recently, we have used these same models to reveal relationships between susceptibility to obesity and motivational response to food cues (Derman and Ferrario, 2018; Alonso-Caraballo and Ferrario, 2019; Derman and Ferrario, 2020; Ferrario, 2020) and inherent differences in the function of mesocorticolimbic systems (Oginsky et al., 2016a; Alonso-Caraballo et al., 2018; Derman and Ferrario, 2018; Vollbrecht et al., 2023) including striatal dopamine (Vollbrecht et al., 2015; Vollbrecht et al., 2016), though see also (Geiger et al., 2008; Lommi et al., 2025), and diet-induced enhancements in nucleus accumbens (NAc) glutamate plasticity (Oginsky et al., 2016b; Oginsky and Ferrario, 2019; Nieto et al., 2023; Fetterly et al., 2024). One advantage to this model is that we know *a priori* which rats are susceptible vs resistant to diet-induced obesity; this allows us to examine inherent differences in brain and behavior without exposure to obesogenic foods. Therefore for clarity, we refer to these lines as obesity-prone (OP) and obesity-resistant (OR) here (and in our prior work) to differentiate studies with and without diet manipulation.

One challenge of selectively bred models is that after initial selection, lines are necessarily maintained in a closed breeding population. This can result in genetic drift and in-breeding (as even an outbred rotational system for establishing new breeding pairs will ultimately fail). To address these concerns, new genes must be introduced into established lines. Here we describe how our lab has maintained and “refreshed” the OP/OR lines originally developed by Levin in 1997. This was achieved by crossing established OP or OR rats with outbred Sprague Dawley rats selected for sensitivity to diet-induced weight gain. We characterized key phenotypes across the selection process and verified weight gain and obesity phenotypes in the final resulting colony. In addition, given the central role of the striatum in motivation for food and opportunistic eating (i.e., eating in the absence of hunger or energy demand), we examined basal striatal function and food motivation in these refreshed lines using whole-cell patch clamping and instrumental procedures.

## Methods

### General Subjects and Conditions

Rats were group housed according to sex with *ad libitum* access to water and food, unless otherwise specified. For all rats bred in house, ages below are exact; for those purchased through commercial vendors ages are approximate. All rats bred in house were weaned at 21 days old.

Established OP and OR colonies (10 breeding pairs of each line) are maintained by the University of Michigan Rodent Breeding Core and their dedicated husbandry staff. The strategy described below was used to ‘refresh’ the colony after we reached the 10^th^ replacement breeders. All procedures were conducted in accordance with The University of Michigan Committee on the Use and Care of Animals in accordance with AAALAC guidelines.

### Initial Screening (F0 generation)

A total of 120 adult (∼45-50 days old on arrival) non-sibling Sprague Dawley rats (60 male, 60 female) were purchased from Charles River Breeding Laboratory (Kingston Facility, New York, USA). Rats were ordered in two separate cohorts of 60 animals, half male and half female. Rats were allowed to acclimate to our housing colony (12h reverse light/dark cycle) for one week. During this time, they had free access to standard lab chow (5LOD; Lab Diet; Indiana, USA; 4 kcal/g: 13.6% fat, 28.9% protein, 57.5% carbohydrates - % of caloric content).

At ∼52-59 days old, all rats were given *ad libitum* access to high-energy diet (D12266B; Research Diets, New Jersey, USA; 4.41 kcal/g: 31.8% fat, 16.8% protein, 51.4% carbohydrates - % of caloric content). This is the same diet composition used to identify and develop the original selectively bred DIO/DR lines (Levin et al., 1997). Body weight was measured approximately one hour after onset of the dark cycle, 3 times per week (MWF) for 2 weeks in males and 3 weeks in females. Males were then rank ordered by total weight gained across the two-week period, whereas females were rank ordered by total weight gained across the three-week period. The top 6 ‘gainers’ and bottom 6 ‘non-gainers’ were identified within each sex, pair housed and placed back on *ad libitum* standard lab chow to await breeding. We then randomly paired the top 6 female gainers with the top 6 male gainers and the bottom 6 female non-gainers with the bottom 6 male non-gainers. The breeding pairs were housed together in ventilated cages for 14 days where they had *ad libitum* access to water and standard lab chow. Females were weighed at the end of 14 days to confirm pregnancy and males were removed at this time. This process was repeated in 2 separate cohorts (60 rats/cohort) and resulted in a total of 24 breeding pairs (N=12 Gainer, N=12 Non-gainer). All F0 breeding pairs exhibited average fecundity (litters sizes of 9-17 pups; 24 litters in total). Representative pups from all 24 litters (F1 generation) were then used to characterize weight gain and related phenotypes, while others were set aside to backcross to existing OP and OR rats (Figure 1 schematic).

**Figure 1:**
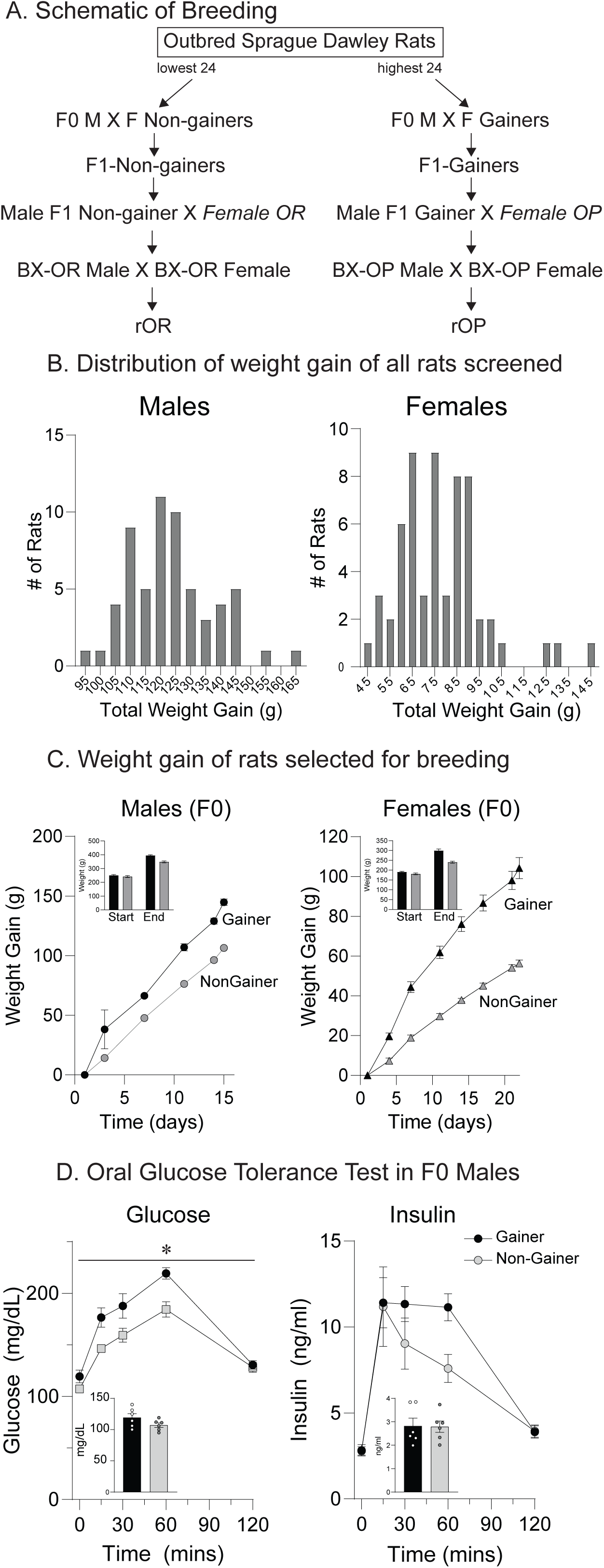
Weight gain and oral glucose tolerance testing in outbred male and female Sprague Dawley rats screened for diet-induced weight gain. A) Schematic of screening and subsequent breeding. Groups in italicized text refer to previously established OP and OR lines. B) Distribution of weight across all outbred rats screened (N=60/sex). C) Cumulative weight gain in rats selected for breeding (N=12/group/sex). The inset shows weight at start and end of high energy diet access for each group. D) Oral glucose tolerance testing in a subset of males after breeding was complete (N=6/group). The insets show baseline glucose (left) and insulin (insulin) prior to oral gavage (2.0g/kg glucose). All data are shown as mean ± SEM.

In addition, 12 of the F0 males used for breeding (∼121 days old; 6 Gainer, 6 Non-gainer) underwent body composition analysis via nuclear magnetic resonance (NMR) and oral glucose tolerance testing (OGTT) after breeding was complete. NMR and OGTT methods are given below.

### F1 generation (offspring of F0 Gainer or F0 Non-gainer pairs) and refreshed OP and OR lines

Two males from each of the 24 F1 litters generated above were set aside to undergo weight-gain, NMR and OGTT studies during adulthood (F1-Gainer N=12; F1-Non-gainer N=12; 65-74 days old at the start of measures). An additional F1 males from each of the 24 litters generated above were paired with OP or OR females from our established colony, with F1s from Gainers paired with OPs, and F1s from Non-gainers paired with ORs (∼ 56-63 days old at pairing) to generate backcrossed offspring (BX-OP, BX-OR). Representative backcrossed male offspring were used for weight-gain, NMR and OGTT studies (48-53 days old; BX-OP N=12; BX-OR N=12). Ultimately, 12 unique, non-sibling breeding pairs were established from backcrossed offspring, constituting the refreshed OP and OR breeding lines that gave rise to refreshed offspring (rOP, rOR).

Subsequent breeding of the rOP and rOR colony followed the Poiley Breeding Method (Bothe, 2010). Both male and female offspring from these refreshed pairs were then used for weight gain, NMR and OGTT studies (76-80 days old at the start; rOP-Male N=20; rOR-Male N=20; rOP-Female N=14; rOR-Female N-14), electrophysiological recordings of striatal neurons (Ns given in Table 1), and to examine instrumental responding for food (51-56 days old at the start of the experiment: rOP-Male N=12; rOR-Male N=12; rOP-Female N=12; rOR-Female N-12). Detailed methods for each procedure and measure follow.

**Table 1:**
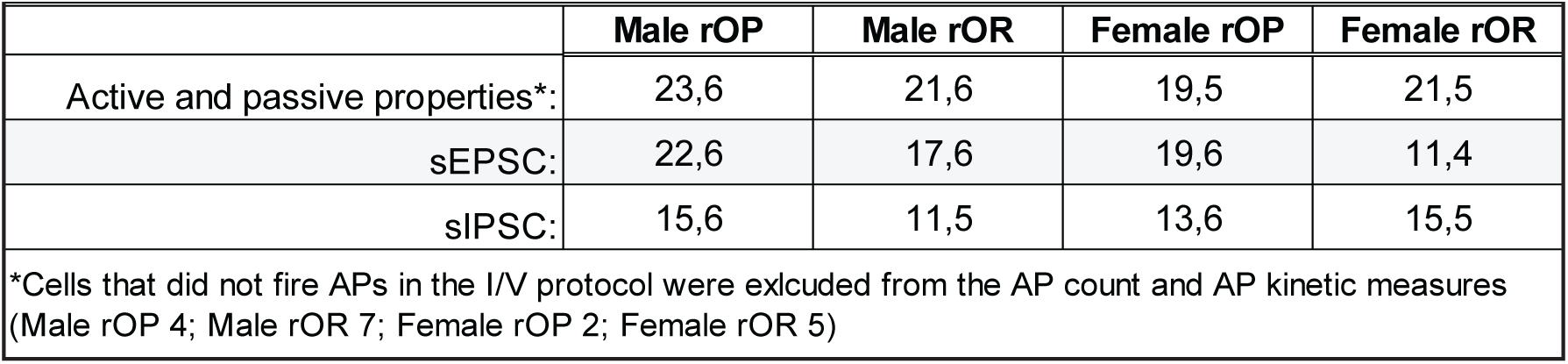
Number of cells and rats per group per recording type given as # of cells, # of rats.

|  |  |  |  |  |
| --- | --- | --- | --- | --- |
|  | Male rOP | Male rOR | Female rOP | Female rOR |
| Active and passive properties*: | 23,6 | 21,6 | 19,5 | 21,5 |
| sEPSC: | 22,6 | 17,6 | 19,6 | 11,4 |
| sIPSC: | 15,6 | 11,5 | 13,6 | 15,5 |
| *Cells that did not fire APs in the I/V protocol were excluded from the AP count and AP kinetic measures (Male rOP 4; Male rOR 7; Female rOP 2; Female rOR 5) |  |  |  |  |

### Weight gain and food intake

Weight gain and food intake studies were conducted when rats reached adulthood (at least 50 days old). All animals were handled once per day for one week prior to diet manipulation. Half the rats were maintained on *ad libitum* standard lab chow while the other half were given *ad libitum* access to a high-energy diet (HED) for 3 weeks. Rats were then weighed 3 times per week (MWF) and home cage food intake was measured at these intervals. The HED and chow diets are closely kilocalorie matched (HED: 4.41 kcal/g, Chow: 4 kcal/g), but the HED contains a larger proportion of fat compared to the chow diet (HED: 31.8% fat. Chow: 13.6% fat).

### Body composition analysis via nuclear magnetic resonance (NMR) and oral glucose tolerance testing (OGTT)

Body composition analysis and OGTT testing were performed by the staff of the University of Michigan Animal Phenotyping Core. Briefly, rats were fasted overnight (∼16h) prior to NMR. Individual rats were placed in a clear plastic holder that was then inserted into an Echo-MRI machine to measure body composition (fat, lean, and free fluid mass). Rats were awake throughout the 2-minute scan. Next, rats were given a 50% glucose solution in 1XPBS via oral gavage (2.0g/kg). Blood was then collected via tail vein bleeding at 0, 15, 30, 60 and 120 minutes after gavage. Blood glucose levels were measured using a glucometer (Acucheck, Roche).

Plasma was isolated via centrifugation (1000 ×g, 4 °C, 10 min) and stored at −20°C for later analysis. Plasma insulin levels were determined using a Millipore rat/mouse insulin ELISA kit. The total area under the curve (AUC) for both glucose and insulin were calculated using the trapezoidal rule (Allison et al., 1995). Glucose-Insulin Index, an indicator of insulin resistance, was calculated as the product of the AUC for glucose and insulin (Henriksen et al., 1998). HOMA-IR index, a homeostatic model assessment of Beta-cell function and insulin resistance was calculated as the product of the fasting glucose and insulin concentration/22.5 (Matthews et al., 1985).

### Whole-cell patch clamping

Established whole-cell patch clamping approaches were used to examine passive and active membrane properties of medium spiny neurons (MSNs) in the NAc core as well as spontaneous excitatory or inhibitory post-synaptic transmission in rOP and rOR rats maintained on standard chow. All recordings were conducted in MSNs from adult rats (70-92 days old) and both sexes were included (See Table 1 for number of cells and rats per recording condition). Slices were prepared as previously described, and slices from females were made when they were in either the metestrus or diestrus phase of their cycle (Alonso-Caraballo et al., 2021; Fetterly et al., 2024). Sectioning was conducted in ice-cold artificial cerebrospinal fluid (aCSF) containing in mM: 125 NaCl, 25 NaHCO_3_, 12.5 glucose, 1.25 NaH_2_PO_4_, 1 ascorbic acid, 3.5 KCl, 0.5 CaCl_2_ and 3 MgCl_2_ (pH = 7.44-7.46 adjusted with NaOH; Osmolality = 295-305 mOsm/kg). The slices were then incubated in aCSF in which KCl and MgCl_2_ were reduced to 2.5 and 1 mM, respectively, while CaCl_2_ was raised to 2.5 mM. Slices remained in aCSF for 20 minutes at 34°C and were allowed to rest for at least 45 minutes at room temperature prior to recording. All solutions were oxygenated throughout (95% O_2_-5% CO_2_).

Patch pipettes were pulled from 1.5 mm borosilicate glass capillaries (Warner Instruments) to resistances of 4-8 MΩ using a horizontal puller (P97, Sutter Instruments). For measures of passive and active properties as well as spontaneous excitatory post-synaptic currents (sEPSCs), pipettes were filled with a solution containing, in mM: 130 K-gluconate, 10 KCl, 1 EGTA, 2 Mg-ATP, 0.6 Na-GTP and 10 HEPES (pH=7.3-7.32 adjusted with KOH; Osmolality = 285-295 mOsm/kg) and recordings were made in current clamp (−200 to 280 pA, 20 pA or 10 pA increments, 500 ms) or voltage clamp mode (−70 mV holding potential), respectively, in aCSF containing 10 µM gabazine. These recordings were Bessel filtered (3 kHZ) and digitized at 10 kHZ. For measures of spontaneous inhibitory post-synaptic currents (sIPSCs), pipettes were filled with a solution containing in mM: 140 CsCl, 2 MgCl_2_, 5 Na-GTP, 10 HEPES, 5 Na-ATP, and 2 QX-314 (pH=7.25 with CsOH; Osmolarity=285-295 mOsm/kg) and recordings were made in voltage clamp mode (−70 mV holding potential) in aCSF containing 20 μM NBQX and 50 μM APV. Electrophysiology data were analyzed using Clampfit 10.7(v10.7, RRID:SCR_011323, Molecular Devices) and Easy Electrophysiology (v2.8.0, RRID: RRID:SCR_021190, Easy Electrophysiology Ltd).

Current-voltage (IV) relationships were determined by calculating the difference between the baseline voltage and the change in voltage 320-420 ms after the start of each current injection. Input resistance was calculated as the slope of the IV curve from −100 to 100 pA. Capacitance (C) was calculated from the τ and input resistance (R; τ = RC). Rheobase was defined as the minimum current injection that elicited an action potential. Measurements of action potential properties were taken from the first action potential elicited. The action potential threshold was determined by the second-derivative method (Sekerli et al., 2004). Voltage depolarization was estimated as the difference between resting membrane potential and threshold voltage. Action potential amplitude was calculated as the difference between action potential peak and voltage threshold. Action potential half-width was defined as the time to reach 50% of the action potential peak. The amplitude of the fast component of the after hyperpolarization (fAHP) was measured within the first 3 ms after reaching the voltage threshold, whereas the medium after hyperpolarization (mAHP) was measured 10 to 50 ms after reaching the voltage threshold. The inter-spike-interval (ISI) was measured in response to the 280 pA current injection and between the peak amplitude of the first two action potentials in the train.

### Instrumental responding for food

Instrumental training and progressive ratio testing were used to evaluate motivation for food. Rats were first given access to food pellets (45mg banana flavored pellets; Bioserv, Product# F0059) overnight in their home cage. They then underwent food cup training during which pellets were delivered on a random interval in the food cup of a standard operant box (Med Associates). Next, rats underwent instrumental training in which pressing the active lever resulted in the delivery of one food pellet and illumination of a cue light above the active lever, 3 sec). Pressing the inactive lever had no consequences, but was recorded. The position of active and inactive levers was left/right counterbalanced relative to the food cup. Rats were given 7 sessions in which each response on the active lever resulted in delivery of a food pellet (fixed ratio 1, FR1, 30 min/session) followed by three progressive ratio (PR) test sessions. During PR testing the number of lever presses required to obtain each subsequent reinforcer increased across the session (5e^(delivery# x 0.2)^-5; adapted from (Richardson and Roberts, 1996; Naleid et al., 2008). During these tests, rats were given two pellets for every ratio completed. The PR session ended automatically when rats did not meet the next ratio requirement within 30 minutes (i.e., breakpoint). Data shown are the average of these three PR sessions.

### Statistics

Data were analyzed in GraphPad Prism 10 using Two-way ANOVAs, Two-way or Three-way Mixed-effects model (REML) or ANOVAs followed by Šídák’s post-tests as appropriate. Statistical analyses and interpretation are informed by guidelines set forth by the American Statistical Association in 2016 (Wasserstein and Lazar, 2016).

## Results

A schematic of the general screening and back-crossing scheme used to refresh existing OP and OR lines is shown in Figure 1A. This approach follows that previously used by Levin and colleagues (personal communication, CRF). Figure 1B shows the frequency distribution of weight gained in all outbred rats that were initially screened, segregated by sex (Male N=60, Female N=60). Figure 1C shows weight gain across time for rats selected for breeding (12 Gainers, and 12 Non-gainers, 6 males and 6 females within each group). Given that rats at the extreme tails of the weight gain distribution were intentionally selected, no statistical analyses were performed. Inset bar graphs show average weight at the start and end of the HED period for the selected rats (2 weeks for males, and 3 weeks for females). These F0 Gainer and F0 Non-gainer groups were then used to generate F1 Gainer and F1 Non-gainer offspring (Figure 1A, B).

After F0 males completed the breeding process (during which they were maintained on standard lab chow), body mass composition was determined followed by a single oral glucose tolerance test (Fat mass: F0 Gainer=6.23% ±0.96; F0 Non-gainer=4.37% ±0.45). Plasma glucose was elevated in F0 Gainer vs F0 Non-gainer males, reached a higher peak in F0 Gainer vs F0 Non-gainer males, and returned to similar levels by the end of the measurement period (Figure 1D, left: Two-way REML main effect of group: F_(1, 10)_=14.1, p=0.004; main effect of time: F(_2.9, 29_)=65.8, p<0.001; group x time interaction: F_(2.9, 29)_=2.5, p=0.08). Although baseline and peak plasma insulin levels were similar between F0 Gainer and Non-gainer groups, plasma insulin remained elevated for longer in F0 Gainers vs Non-gainers (Figure 1D, right: Two-way REML main effect of line: F_(1, 10)_=1.14, p=0.03; main effect of time: F_(1.29, 12.9)_=39.7, p<0.001; line x time interaction: F_(1.29, 12.9)_=1.7, p=0.21). Thus, elevations in weight, fat mass, and plasma glucose were accompanied by a sustained insulin response in F0 Gainers compared to F0 Non-gainers.

Figure 2 shows weight gain and food intake of male F1 Gainer and F1 Non-gainer offspring (panels A,B), and male offspring resulting from backcrossing F1 Gainer and F1 Non-gainers to existing OP (BX-OP) and OR (BX-OR) rats, respectively (panels C,D). As expected, weight gain was greater in F1 Gainers vs F1 Non-gainers maintained on chow, and this difference was further amplified by *ad libitum* access to HED (Figure 2A: Three-way REML main effect of line: F_(1, 20)_=5.98, p=0.02; main effect of diet: F_(1, 20)_=11.1, p=0.003; line x diet and line x diet x time interaction, P=0.72-0.94). Food intake was also greater in F1 Gainers vs F1 Non-gainers, regardless of the diet available (Figure 2B: Three-way REML main effect of line: F_(1, 20)_=18.8, p=0.0003, diet x line interaction, p=0.22). In addition, both groups tended to eat less HED than chow (Figure 2B: Three-way REML main effect of diet: F_(1, 20)_=4.6, p=0.04), though this difference was more pronounced in F1 Gainers vs F1 Non-gainers (Figure 2B: open vs closed circles). Kilocalorie per gram is closely matched between HED and chow diets (HED: 4.41 kcal/g, Chow: 4 kcal/g), but the HED contains a larger proportion of fat compared to the chow diet (HED: 31.8% fat. Chow: 13.6% fat). This promotes weight gain, particularly in susceptible populations, even when consumption of HED is similar to that of chow.

**Figure 2:**
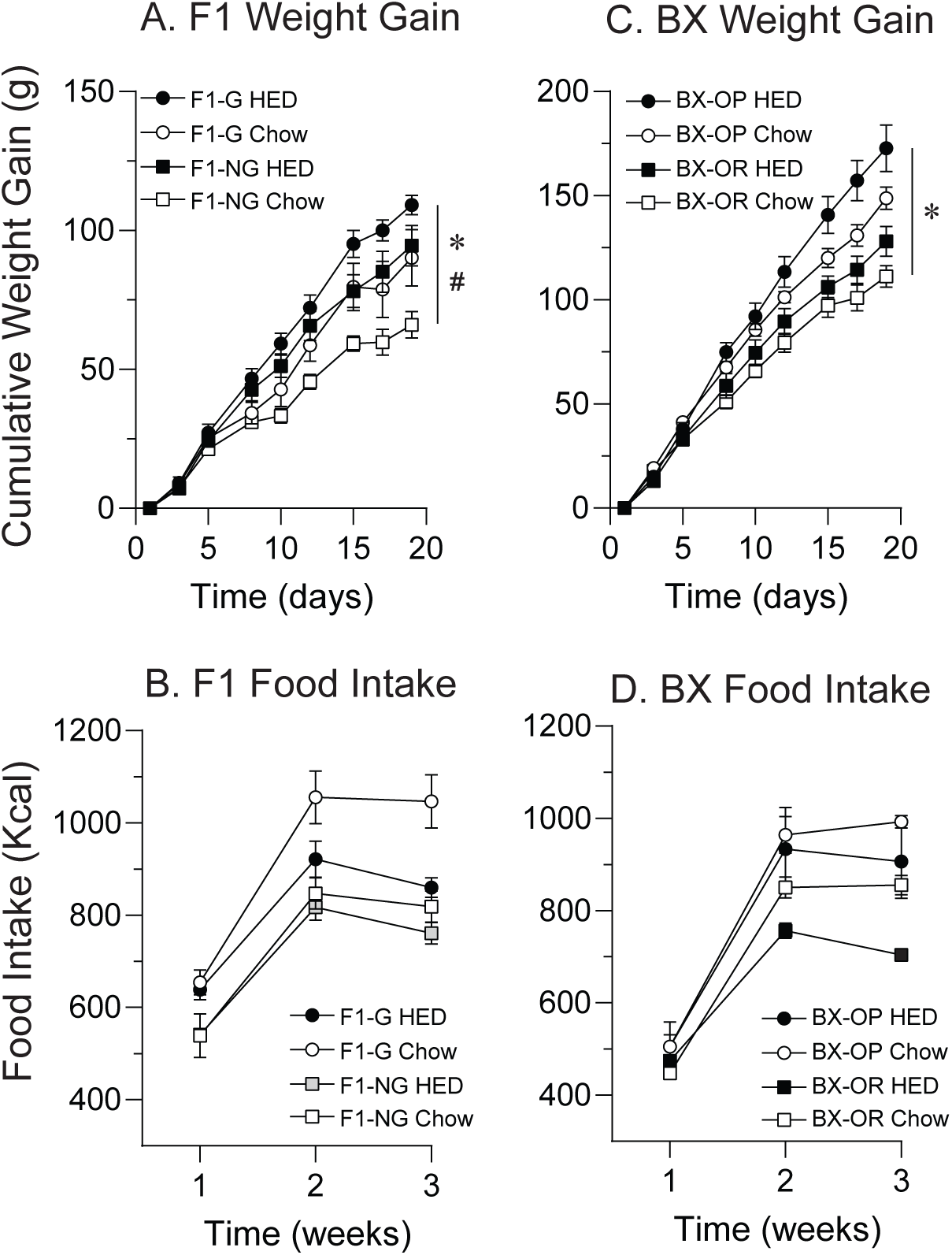
Weight gain and food intake in male F1 (A,B) and backcrossed (C, D) male rats given *ad libitum* access to standard lab chow or high energy diet (HED; N=12/group); *= Three-way REML main effect of line, p<0.01; #= Three-way RELM main effect of diet, p<0.01; $=Three-way REML time x diet and time x line interaction, p<0.01.

The weight gain phenotypes observed in the F1 generation were more exaggerated in BX-OP vs BX-OR groups (Figure 2C,D). Specifically, differences in weight gain between BX-OP and BX-OR groups were generally more pronounced (Figure 2C: Three-way REML main effect of line: F_(1, 20)_=20.4, p=0.0002; line x diet and line x diet x time interaction, P=0.72-0.38) and HED produced greater weight gain than chow in both groups, with the greatest gain in BX-OPs compared to BX-ORs (Figure 2C: Three-way REML main effect of diet: F_(1, 20)_=3.94, p=0.06; time x diet interaction: F_(1.7, 34.6)_=8.58, p=0.001; time x line interaction: F_(1.7, 34.6)_=22.55, p<0.001). Food intake was greater in BX-OPs compared to BX-ORs, regardless of the diet available (Figure 2D: main effect of line: F_(1, 20)_=8.86, p=0.0075; main effect of diet: F_(1, 20)_=1.93, p=0.18; diet x line interaction: F_(1, 20)_=0.17, p=0.68).

Figure 3 shows the effects of HED on body composition, glucose, and insulin levels in BX-OP and BX-OR male offspring. BX-OP were heavier than BX-OR, regardless of diet (Figure 3A: Two-way ANOVA; main effect of line: F_(1, 16)_=77.33, p<0.0001; no main effect of diet p=0.36; line x diet interaction: F_(1, 16)_=4.10, p=0.05). Moreover, examination of body composition revealed that fat mass was greater in BX-OP vs BX-OR offspring, and that this difference was amplified by HED consumption (Figure 3B: Two-way ANOVA; main effect of line: F_(1, 16)_=37.09, p<0.0001; main effect of diet: F_(1, 16)_=24.46, p<0.0001; no line x diet interaction p=0.21). Consistent with this, lean mass was also reduced to a greater extent by HED consumption in BX-OPs compared to both BX-OR or BX-OP offspring eating chow (Figure 3C: Two-way ANOVA; main effect of line: F_(1, 16)_=26.91, p<0.0001; main effect of diet: F_(1, 16)_=15.47, p<0.001; no line x diet interaction p=0.45). After an overnight fast, all rats were given an oral glucose tolerance test. Figure 3D,E show fasted concentrations of blood glucose and plasma insulin (respectively), and panel F shows the HOMA-IR, an estimate of insulin resistance. Fasted glucose and insulin levels were greater in BX-OP vs BX-OR offspring, with an expected exacerbation due to HED consumption (Figure 3D: Two-way ANOVA; main effect of line: F_(1, 16)_=4.93, p<0.04; main effect of diet: F_(1, 16)_=9.154, p<0.008; no line x diet interaction p=0.43; Figure 3E: main effect of line: F_(1, 16)_=18.75, p<0.0005; main effect of diet: F_(1, 16)_=11.01, p<0.004; no line x diet interaction p=0.56). This resulted in a higher HOMA-IR ratio in BX-OP vs BX-OR offspring given chow that was further elevated following HED consumption (Figure 3F: Two-way ANOVA main effect of line: F_(1, 16)_=6.19, p<0.02; main effect of diet: F_(1, 16)_=22.38, p<0.0002; line x diet interaction: F_(1, 16)_=6.19, p=0.02). HED-induced slowed glucose clearance in both groups, with larger effects in BX-OP vs BX-OR offspring (Figure 3G: Three way REML main effect of line: F_(1, 16)_=28.68, p<0.0001; main effect of diet: F_(1, 16)_=12.31, p=0.0029; time x line interaction: F_(4, 64)_=5.184, p=0.001; time x diet interaction: F_(4, 64)_ =3.38, p=0.01; line x diet interaction: F_(1, 16)_=4.43, p=0.05; no time x diet x line interaction, p =0.12; Two-way REML: main effect of line BX-OP-Chow vs BX-OR-Chow: F_(1, 8)_ = 11.3, p=0.01; main effect of line BX-OP-HED vs BX-OR-HED: F_(1, 8)_ = 18.16, p=0.03). This was accompanied by slight elevations in baseline insulin, corresponding enhancements in peak insulin and a slower decline in insulin for rats given HED, and in BX-OP vs BX-OR offspring (Figure 3H: Three way REML main effect of line: F_(1, 16)_=21.20, p<0.0003; main effect of diet: F_(1, 16)_=8.62, p=0.01; time x diet interaction: F_(2.683, 42.93)_=4.81, p=0.007; no line x diet interaction, p=0.84; no time x line interaction, p=0.455; no time x diet x line interaction, p =0.36; Two-way REML: main effect of line BX-OP-Chow vs BX-OR-Chow: F (1, 8) = 28.06, p<0.0007; main effect of line BX-OP-HED vs BX-OR-HED: F_(1, 8)_ = 6.117 p=0.04). Overall, BX-OP offspring maintained on chow were heavier, had more fat mass and were both hyperglycemic and hyper insulinemic compared to BX-ORs. These differences were further exacerbated by prolonged consumption of HED.

**Figure 3:**
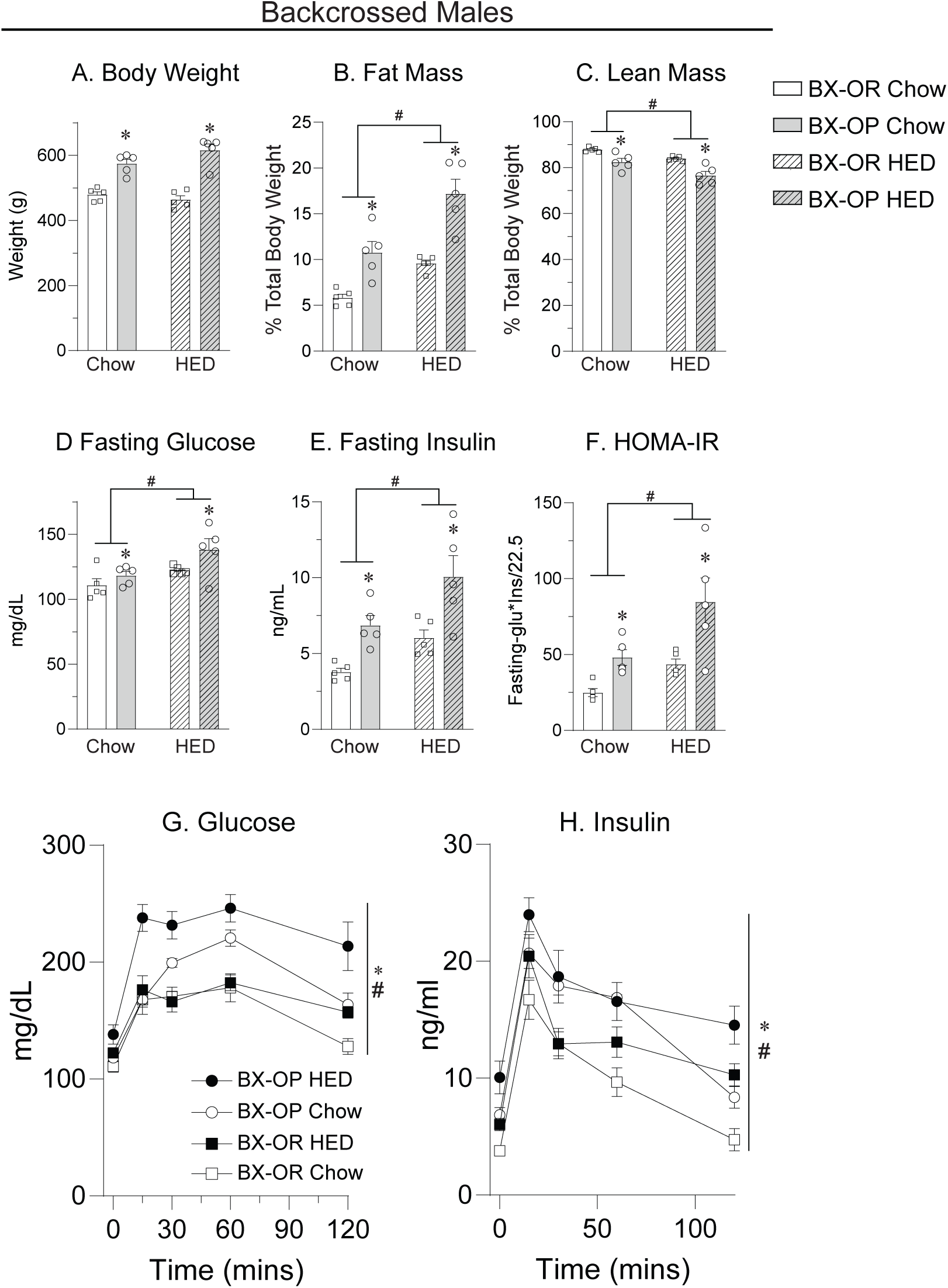
Effects of high energy diet (HED) on body composition, glycemia and the insulin response in male offspring from backcrossed (BX) OP and OR lines (N=12/group). Body weight (A), fat mass (B), lean mass (C), fasted blood glucose (D), fasted plasma insulin (E), Homeostatic Model Assessment of Insulin Resistance (HOMA-IR) ratio (F), and glucose (G) and insulin (H) following oral glucose tolerance testing in backcrossed OP (BX-OP) and OR (BX-OR) males given chow or HED *ad libitum*. *= Three-way REML main effect of line; # Three-way REML main effect of diet, p < 0.05.

Figure 4 shows effects of HED on weight gain and food intake from male and female offspring of the refreshed OP and OR lines (rOP, rOR). As expected, weight gain was greater in rOP vs rORs, and HED resulted in more pronounced weight gain in males (Figure 4A: Three-way REML, main effect of line: F_(1, 36)_=16.43, p=0.0003; main effect of diet: F_(1, 36)_=23.04, p<0.0001; line x diet x time interaction: F_(1.134, 40.31)_=6.251, p=0.01) and in female rOPs compared to rORs (Figure 4B: Three-way REML, main effect of line: F_(1, 24)_=17.43, p=0.0003; main effect of diet: F_(1, 24)_=7.0, p=0.01; no line x diet or line x diet x time interaction, P=0.19-0.31). By the end of the HED consumption period average weight (±SEM) in grams for males was as follows: rOP-Chow 386.5 ±58.2; rOR-Chow 368.9 ±50.26; rOP-HED 405.2 ±78.9; rOR-HED 371.5 ±55.66. By the end of the HED consumption period weight in grams for females was as follows: rOP-Chow 247.1 ±27.43; rOR-Chow 224.6 ±21.79; rOP-HED 256.6 ±42.57; rOR-HED 232.9 ±30.5). In addition, rOP males ate more than rORs and this difference became more pronounced across time in HED groups (Figure 4C: Three-way REML, main effect of line: F_(1, 36)_=10.23, p=0.003; time x diet interaction: F_(1.437, 51.75)_=8.22, p=0.002; Sidak’s posttest, rOR-HED vs rOP-HED, p<0.05 week 2 & 3). A similar pattern was observed in females (Figure 4D: Three-way RELM, main effect of line: F_(1, 24)_=5.02, p=0.03; time x diet interaction: F_(1.551, 37.22)_=7.82, p=0.003).

**Figure 4:**
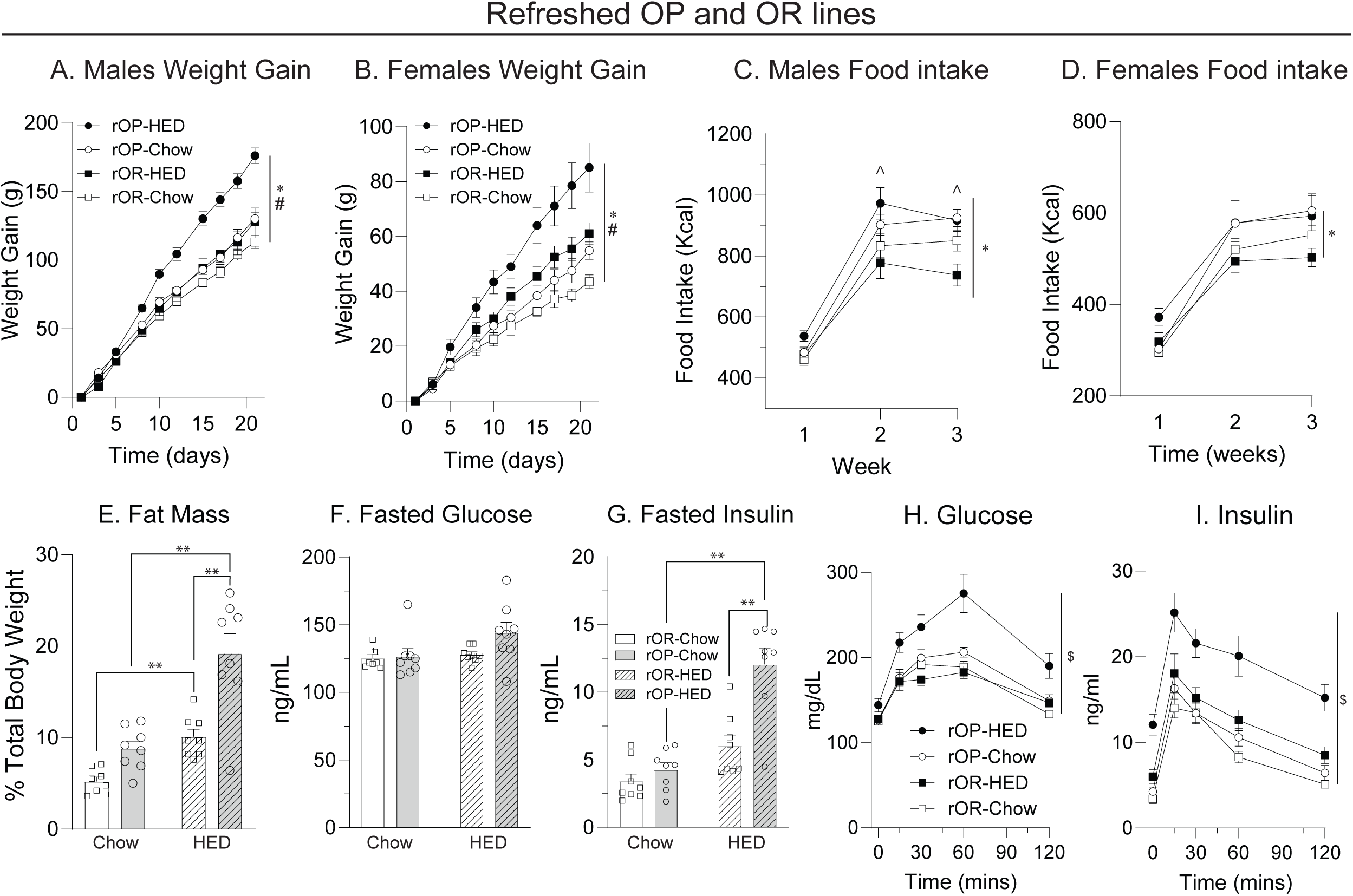
Effects of high energy diet (HED) on weight and food intake in male and female offspring of the refreshed OP and OR lines (rOP-Male N=20; rOR-Male N=20; rOP-Female N=14; rOR-Female N-14). Weight gain in males (A) and females (B). Food intake in males (C) and females (D). Fat mass (E), fasted blood glucose (F), and fasted plasma insulin (G) prior to oral glucose tolerance testing in males. Glucose (H) and insulin (I) following oral glucose tolerance testing. *=Three-way REML main effect of line; #= Three-way REML main effect of diet, p < 0.05, ^=Sidak’s post-test rOP-HED vs rOR-HED, p<0.05; **= Sidak’s post-test, p<0.02 between pairs as indicated; $=Two-way ANOVA main effect rOP-Chow vs rOP-HED and rOP-HED vs rOR-HED.

A separate group of refreshed male offspring were given *ad libitum* access to chow or HED, and fasted glucose, insulin and the response to an oral glucose tolerance test were evaluated. Average weight (± SEM) for these rats was: rOR-Chow 408 ±13.1; rOP-Chow 446.3 ±14.1; rOR-HED 422.5 ±16.17; rOP-HED 511.7 ±20.1). As expected, HED resulted in increased fat mass in both lines, but this effect was larger in rOP-HED vs rOR-HED offspring (Figure 4E; Two-way REML main effect of line: F_(1, 28)_=24.82, p<0.0001; main effect of diet: F_(1, 28)_=36.21, p<0.0001; line x diet interaction: F_(1, 28)_=4.625, p=0.04; Sidak’s post-test: rOP-Chow vs rOP-HED, p<0.001; rOR-Chow vs rOR-HED, p<0.02; rOP-Chow vs rOR-Chow p=0.11; rOP-HED vs rOR-HED, p<0.0001). Basal fasted glucose level did not differ substantially between groups, but was highest in rOPs maintained on HED (Figure 4F; Two-way ANOVA; main effect of diet F_(1, 28)_=3.890, p=0.06; main effect of line: F_(1, 28)_ = 2.905, p=0.10; diet x line interaction F_(1, 28)_=2.134, p=0.155). Furthermore, basal fasted insulin was similar between rOP- and rOR-Chow groups, while HED resulted in greater fasted insulin levels in rOPs (Figure 4G; Two-way ANOVA significant line x diet interaction F_(1, 28)_=9.9, p=0.004, Sidak’s post-test rOP-Chow vs rOP-HED, p<0.001; rOR-Chow vs rOR-HED, p=0.065; rOP-HED vs rOR-HED, p<0.0001). Glucose clearance (Figure 4H) was generally similar between rOP-chow, rOR-chow, and rOR-HED groups, but blunted in rOPs maintained on HED compared to rOP-Chow and rOR-HED groups (Figure 4H: Three-way REML main effect of line: F_(1, 28_ =17.28, p=0.0003; time x line interaction: F_(2.966, 83.05)_=6.147, p=0.0008; time x line x diet interaction F_(2.966, 83.05)_=2.972, p=0.04; Two-way RM ANOVA rOP-Chow vs rOP-HED main effect of diet F_(1, 14)_= 9.14, p=0.009; rOP-HED vs rOR-HED main effect of diet F_(1, 14)_=14.20, p=0.002). Consistent with this, the insulin response was similar in chow groups, and enhanced by HED in rOP and rOR rats, with larger effects in rOPs (Figure 4I: Three-way RELM, main effect of line: F (1, 28) = 13.88, p=0.001; main effect of diet: F_(1, 28)_=29.77, p<0.0001; line x diet interaction F_(1, 28)_=6.198, p=0.02; no line x diet x time interaction, p=0.95; Two-way RM REML rOP-Chow vs rOP-HED main effect of diet F_(1, 14)_=22.1, p=0.0003; no time x diet interaction, p=.85; rOP-HED vs rOR-HED main effect of diet F_(1, 14)_=12.6, p=0.003; no time x diet interaction, p=0.93).

We also examined intrinsic membrane and firing properties as well as synaptic transmission (excitatory and inhibitory) in MSNs of the NAc core from male and female rOPs and rORs maintained on standard lab chow (Ns given in Table 1). Overall, we did not find robust differences between the rOP and rOR lines. However, we did find interesting sex differences in MSN membrane properties and excitatory transmission (Figure 5), and some of these sex differences were more pronounced in rOPs than rORs. Specifically, there were larger changes in membrane potential in response to positive current injection in females compared to males (Figure 5A; Three-way REML, main effect of sex F_(1,80)_=11.09, p<0.01 and sex x current injection F_(24,1920)_=7.60, p<0.0001; no main effect of line, line x current injection, or sex x line x current injection interaction, P=0.90-0.99). This was not due to differences in rheobase (Figure 5B) or resting membrane potential (data not shown). However, MSN capacitance was lower in females than males (Figure 5C; Two-way REML main effect of sex, F_(1,80)_=15.05, p<0.001; no main effect of line or line x sex interaction, P=0.26-0.65), and input resistance was higher in females than males (data not shown: Two-way REML main effect of sex, F_(1,80)_=6.02, p<0.05; no main effect of line or line x sex interaction, P=0.53-0.83). These data indicate that cell size was smaller in females than males, consistent with the greater depolarization in response to current injection found in females compared to males (Figure 5A).

**Figure 5:**
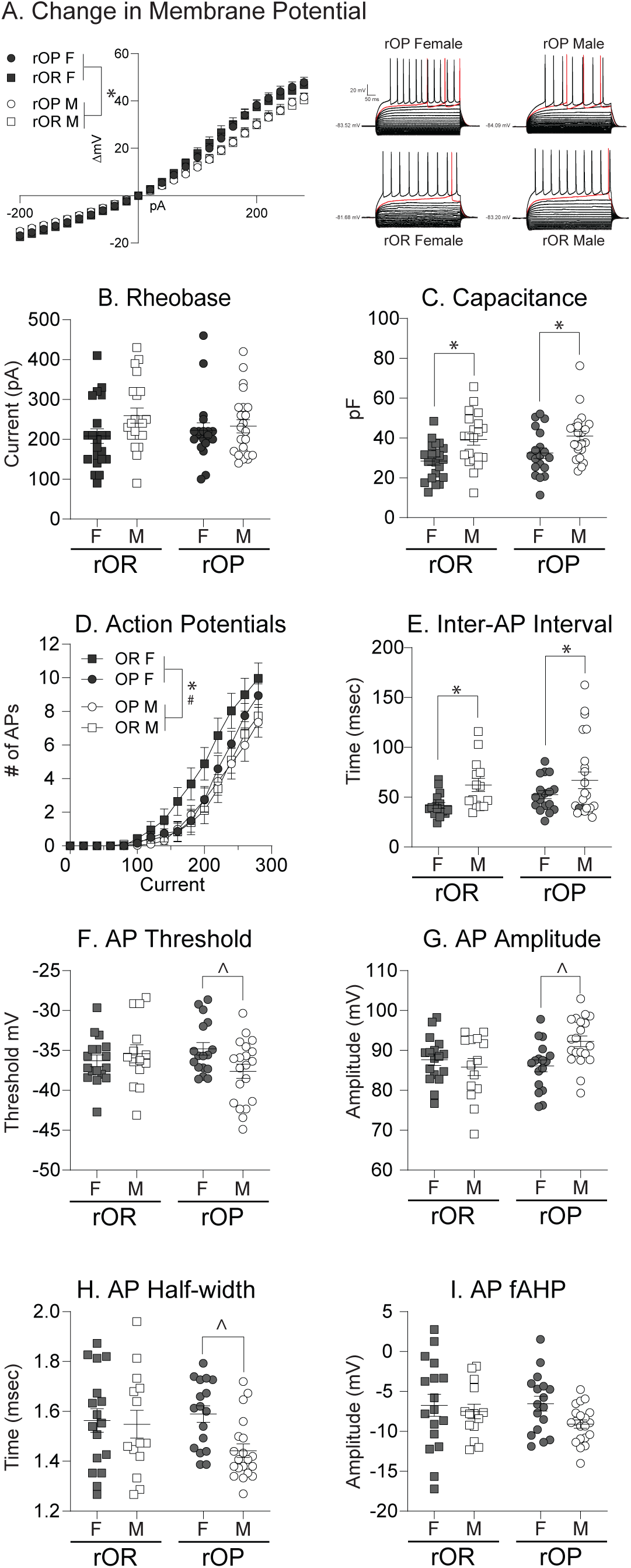
Examination of intrinsic membrane and firing properties of NAc core medium spiny neurons from rOP and rOR rats reveals sex differences that are largely similar across rOP and rOR lines (see Table 1 for Ns). Example traces are shown in panel A. For all panels, data from females are shown in closed symbols, and data from males are shown in open symbols. Current/voltage (IV) relationships (A), rheobase (B), cell capacitance (C), the number of action potentials (AP) across current injection (D), the inter-action potential interval (E), and action potential threshold (F), amplitude (G), half-width (H) and fast after hyperpolarization (I; fAHP). *=Three-way REML or Two-way ANOVA main effect of sex, p<0.05; #=Three-way REML or Two-way ANOVA sex x line interaction, p<0.05; ^=Two-way ANOVA sex x line interaction, Sidak’s post-test, p<0.05.

Sex differences were also present in active membrane properties and aspects of the action potential (Figure 5D-I). Specifically, the number of action potentials elicited by increasing current injection was greater in females compared to males (Figure 5D; Three-way ANOVA, main effect of sex: F_(1,64)_=4.17, p=0.045; sex x current injection interaction: F_(1442,92.31)_=3.93, p<0.05). Consistent with this, the inter-spike interval at 280 pA was shorter in females compared to males, again in both rOP and rOR groups (Figure 5E: Two-way ANOVA main effect of sex: F_(1,66)_=6.87, p=0.01; no sex x line interaction, p=0.64). Sex differences in other aspects of the action potential, including the firing threshold, peak amplitude of the first action potential, and the half-width of this response (Figure 5F-H) largely lay within the OP group (Two-way ANOVAs, threshold: significant sex x line interaction F_(1,64)_=4.32, p=0.04, Sidak’s male vs female OP p=0.04; male vs female OR p=0.75; peak amplitude: significant sex x line interaction F_(1,64)_=6.46, p=0.01, Sidak’s male vs female OP p=0.01; male vs female OR p=0.68; half-width: significant main effect of sex, F_(1,64)_=4.02, p=0.049, Sidak’s male vs female OP p=0.02; male vs female OR p=0.96). Overall, sex differences in action potential firing, while present, are less consistent across measures and more variable across phenotype than sex differences in intrinsic membrane properties.

When synaptic transmission was examined, we did not find any differences in spontaneous inhibitory post-synaptic transmission (data not shown) or differences in excitatory transmission between rOP and rOR lines (Figure 6, example traces are shown in panel B inset). However, we again observed sex differences with reduced frequency of sEPSCs in females vs males (Figure 6A, Two-way ANOVA main effect of sex, F_(1,65)_=4.866, p<0.05, no main effect of line, or sex x line interaction, P=0.74-0.9) and a significant rightward shift in the cumulative frequency distribution such that females had more events with a longer inter-event intervals (Figure 6B, Three-Way REML, significant sex x inter-event interval interaction, F_(50,2952)_=4.49, p<0.0001; no main effect of line or sex x line interaction, P=0.98-0.99). In addition, the average amplitude of sEPSCs was larger in females than males (Figure 6C; Two-way ANOVA main effect of sex, F_(1,52)_=8.97, p<0.01; no main effect of line or sex x line interaction, P=0.84-0.96). In addition, there was a small but significant shift in the amplitude distribution such that in females more events fell into and slightly above the peak amplitude range (Figure D; Three-Way REML significant sex x amplitude interaction, F_(10,512)_=3.00, p<0.01). Finally, sEPSC decay was similar across groups and sex (data not shown). Thus, overall, the frequency of sEPSCs was less in females than males, but these events tended to be of larger amplitude.

**Figure 6:**
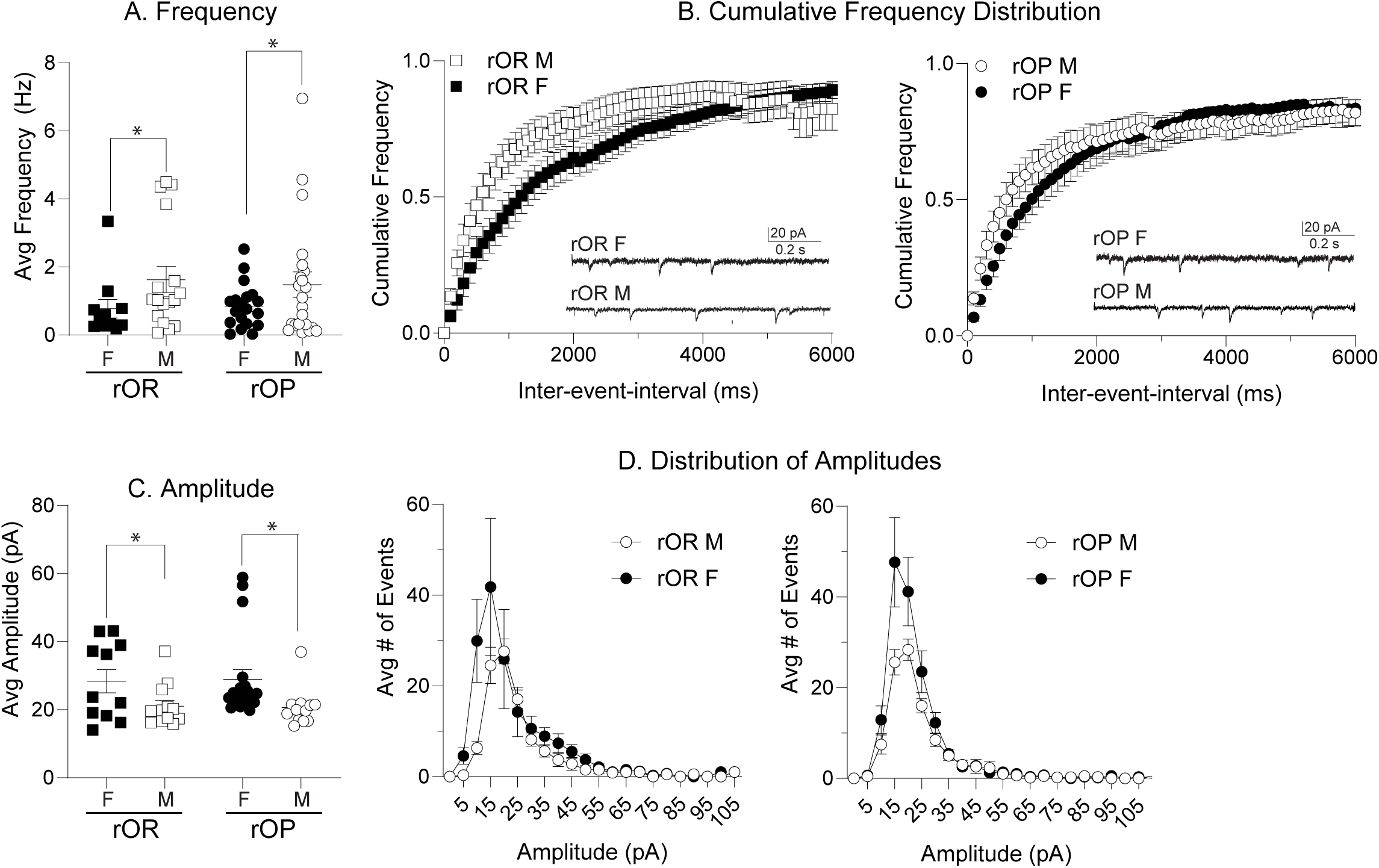
Sex differences in the frequency and amplitude of spontaneous excitatory post-synaptic currents (sEPSCs) in the NAc core of rOP and rOR rats (see Table 1 for Ns). For all panels, data from females are shown in closed symbols, and data from males are shown in open symbols. sEPSC Frequency (A), and the cumulative frequency distribution (B) for rOR (left) and rOP (right) groups. Panel B insets shows example sEPSC traces. sEPSC Amplitude (C) and distribution of amplitudes (D) for rOR (left) and rOP (right) groups. *=Two-way ANOVA main effect of sex, p<0.05; see also results.

Finally, we measured motivation for food in male and female rOP and rORs maintained on standard lab chow using instrumental procedures (N=12 per sex per group). Sex was included as a factor in our statistical analyses, but data are graphed together because we did not find any effects of sex. Figure 7A shows initial acquisition in rOP and rORs, with data from males and females segregated in the inset. The rate of acquisition was similar between rOP and rOR groups, with discrimination between active and inactive levers emerging at similar rates within the first FR1 training session for both groups (Figure 7B, Session 1; Three-way REML main effect of lever: F_(1, 92)_ = 15.46, p=0.0002; time x lever interaction: F_(2.861, 263.2)_ = 11.23, p<0.0001; no main effect of line, line x time, line x lever, or line x time x lever interaction: P=0.65-0.98). However, the magnitude of active lever responding during the later phase of training was greater in rOP vs rOR groups (Figure 7A: Three-way REML session x line interaction: F_(3.195, 140.6)_ = 3.049, p=0.03; no main effect of sex, line or interactions between factors, P=0.31-0.99). This enhancement in responding for food occurred at the start of each session beginning at session 4 (Figure 7B). Rats next underwent progressive ratio testing, where the number of responses required to receive a single food pellet increased exponentially across the session. Figure 7C shows the average break point (i.e., the last ratio completed) across the three PR test sessions, and Figure 7D shows the average time it took for rats to reach their break point (data from males are shown in open symbols and data from females are shown in closed symbols for both panels). Break point did not differ between rOPs and rORs. However, rOPs reached their break point more quickly than rORs (Figure 7D: Two-way ANOVA: main effect of line: F_(1, 44_ =4.268, P=0.04; no main effect of sex p=0.12 or sex x line interaction, p=0.44).\ Overall, motivation to work for food was greater in rOP vs rORs of both sexes. This difference was apparent when work requirements were low and led rOPs to reach their break points faster than rORs.

**Figure 7:**
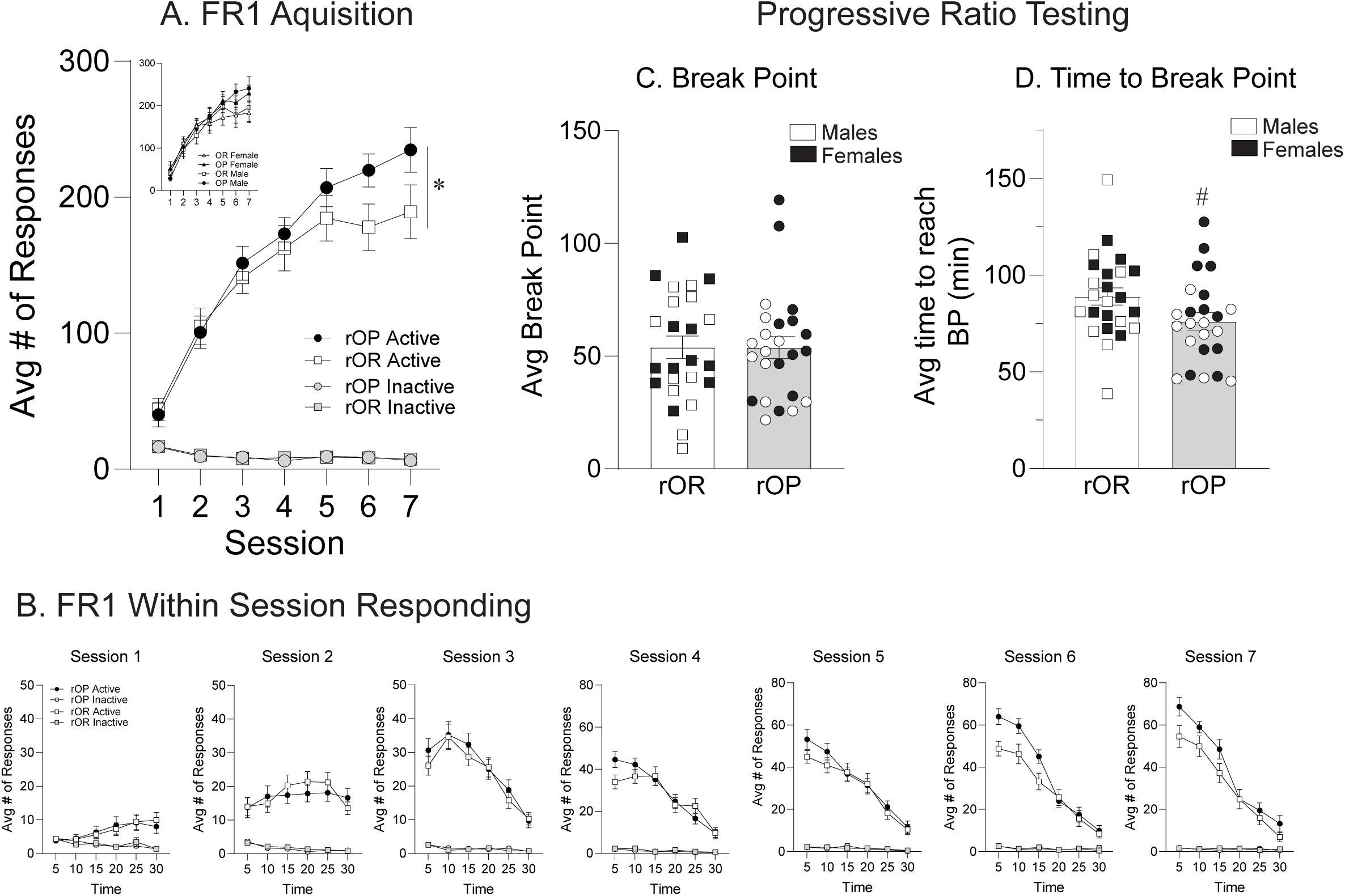
Motivation to obtain food is greater in rOP vs rOR rats (N=12/group/sex). A) Initial acquisition of instrumental responding for food (fixed ratio 1, FR1). The inset shows these same data segregated by sex. The rate of acquisition is similar across groups, and the magnitude of responding is higher in rOP vs rORs. B) Within session responding during each FR1 session. rOPs and rORs Discriminate between the active and inactive levers by the end of the first session, and enhancements in active lever responding in rOPs emerge early in session 4 and are maintained through session 7. C) Average break point reached during progressive ratio testing with data from females in closed symbols and data from males in open symbols. D) Average time to reach break point (BP) during progressive ratio testing with data from females in closed symbols and data from males in open symbols. rOPs Reached their break point faster than rORs, consistent with greater motivation for food. *=Three-way REML session x line interaction, p<0.05; #=Two-way ANOVA main effect of line, p<0.05.

## Discussion

Data from human and rodent studies show that risk of developing obesity is enhanced by genetic factors that interact with the food environment in ways that slow metabolism, alter motivation, and promote unhealthy eating patterns. Rodent models of susceptibility and resistance to diet-induced obesity, including the selectively bred OP and OR lines described here, have been invaluable for understanding these interactions and their consequences on physiology and behavior. Maintaining selectively bred lines in closed populations can lead to genetic drift and potential inbreeding, necessitating the periodic introduction of genes from outside the colony. Here we describe a strategy for refreshing the OP and OR lines, characterize key phenotypes across this process, and verify that the resulting “refreshed” colonies retained key metabolic and behavioral characteristics of these models.

Outbred Sprague Dawley rats were purchased from the Charles River Breeding Laboratory’s Kingston Facility in New York, as this vendor and location have been used as the source of outbred rats since the initial creation of this line (personal communication, BE Levin), and breeding facilities have their own standards of practice that can influence biology and behavior even within the same rat strain (e.g., Gileta et al., 2022). We screened these F0 breeders for propensity to diet-induced weight gain and then bred the top 12 female and male gainers to each other and the bottom 12 female and male Non-gainers to each other to generate an F1 generation that was then back-crossed to our existing OP and OR lines (see schematic in Figure 1A). We chose this approach, rather than using the F0 rats themselves as founders for backcrossing, to provide some amplification of the selected trait and to minimize introducing potential effects of HED consumption into our breeding colony. For example, 2 weeks of HED consumption was sufficient to increase fasted plasma glucose and prolong increases in insulin during the oral glucose tolerance test in male gainers compared to Non-gainers (Figure 1D). Additionally, potential epigenetic responses to HED could be important to phenotypic differences. Consistent with this, epigenetic modifications contribute to obesity susceptibility and are likely an inherent feature of the OP/OR models (Gorski et al., 2007; Bouret et al., 2015).

As expected, phenotypic differences were modest but present in F1 Gainer vs Non-gainer offspring (Figure 2A,B), more pronounced following backcrossing to existing lines (Figure 2C,D and Figure 3), and robust in the final refreshed colony (rOP vs rOR, Figure 4). Importantly, the rOP and rOR lines recapitulated core phenotypes of the original model; rOPs showed greater weight gain while maintained on chow than rORs and this was exacerbated by consumption of HED in both sexes (Figure 4A,B). This was also accompanied by greater fat mass, hyperinsulinemia, and blunting of glucose tolerance in rOP males maintained on chow vs HED (Figure 4E-I). Conversely, rOR males given HED were more similar to their rOR chow counterparts in weight gain, body composition and metabolic profile, and showed less robust effects of HED compared to rOPs. These overall patterns are consistent with prior characterizations of OP/DIO and OR/DR phenotypes (e.g., Levin et al., 1997; Madsen et al., 2010; Vollbrecht et al., 2015; Alonso-Caraballo et al., 2018) and highlight the utility of these lines for isolating effects of obesogenic foods from weight gain and metabolic dysfunction. Furthermore, weight gain trajectories and metabolic profiles do not differ dramatically in rOP and rOR males maintained on chow particularly during young adulthood (Figure 4 and Vollbrecht et al., 2015; Alonso-Caraballo et al., 2018). Thus, this model is also particularly useful for examining inherent neurobehavioral differences associated with susceptibility to weight gain prior to the development of obesity and without the introduction of an overt diet manipulation.

We examined intrinsic cell properties, action potential properties (Figure 5), and synaptic transmission (Figure 6) in MSNs from the NAc core of male and female rOP and rOR groups because of the central role of the NAc in food-seeking and motivation as well as established differences in diet-induced NAc plasticity in these models (Oginsky et al., 2016b; Oginsky and Ferrario, 2019; Nieto et al., 2023; Fetterly et al., 2024). We did not find robust differences in MSN physiology between the rOP and rOR lines. However, we did find interesting sex differences in MSN membrane properties, firing, and excitatory transmission, and some of these sex differences were more pronounced in rOPs than rORs. First, there were larger changes in membrane potential in response to positive current injection in females compared to males. This did not appear to be due to shifts in membrane excitability per se, but rather smaller cell size in females than males (indicated by lower capacitance; Figure 5C). Second, action potentials in females were more frequent and had a shorter inter-spike interval in response to the same current injection compared to males (Figure 5D,E). There were also sex differences in other aspects of the action potential (firing threshold, peak amplitude and half-width; Figure 5F-H) that were more apparent within OPs. Thus, while present, sex differences in the action potential itself were more variable across measures and phenotype than sex differences in intrinsic membrane properties. Finally, regarding synaptic transmission, we found that while spontaneous excitatory events were less frequent in females than males, they tended to be of larger amplitude (reflected by a shift in the amplitude distribution and average peak). This pattern is suggestive of greater postsynaptic AMPA receptor number, and less excitatory synaptic input in females than males, and is consistent with evidence for greater excitatory synaptic strength in female than male NAc, although this literature is mixed and underdeveloped (Kniffin and Briand, 2024).

We previously found enhancements in intrinsic excitability of NAc core MSNs in male and female OPs compared to ORs maintained on chow, lower GluA1 surface expression, and reduced AMPA/NMDA ratio in OP vs OR groups (Oginsky et al., 2016a; Derman and Ferrario, 2018; Oginsky and Ferrario, 2019; Alonso-Caraballo et al., 2021). No differences in intrinsic properties or sEPSCs were found between rOP vs. rORs here (Figure 6). Prior results were reproducible across several years and different experimenters, thus we do not think they were spurious. However, it’s possible that the refreshing approach diluted these neural phenotypes, and that replacement of breeding pairs across time may amplify phenotypes. Additionally, recordings here were conducted in rats spanning a relatively wide age range (71-92 days old), whereas in our previously published studies rats were more similar in age within cohort (∼1 week). Thus, age could be an additional contributing variable.

We also examined instrumental responding for food using a fixed ratio 1 (FR1) schedule of reinforcement and progressive ratio testing (Figure 7). Like previous results from these lines (Vollbrecht et al., 2016; Fetterly et al., 2024), rOP and rOR rats learned this task similarly, discriminating between active and inactive levers by the end of the first 30-minute session (Figure 7B), but rOPs maintained a higher rate of responding compared to rORs in later sessions. This difference emerged during the 4^th^ session and became more pronounced across sessions 5 through 7, with rOPs responding much more during the first half of each session than ORs and maintaining a similar level of responding to rORs for the remainder of the session. This carried over to progressive ratio testing, where although break points were similar between groups, rates of responding were greater in rOPs, such that they reached their breakpoint more quickly (Figure 7C,D). Thus, motivation for food was greater in rOPs vs rORs maintained on standard lab chow, and this difference was more pronounced when work requirements were low. This is consistent with differences in food intake, the general phenotypes of these lines, and with inherent differences in the function of mesocorticolimbic circuits that underlie motivation.

In summary, results here show that the strategy described above results in refreshed OP and OR lines retain the core metabolic and behavioral characteristics of the original models. This provides a well-characterized colony for continued investigation of neurobehavioral factors that promote susceptibility to obesity and diet-induced neurobehavioral plasticity in the context of health and weight gain. Rodent models like these are necessary to the development of obesity treatment and prevention strategies.

## Acknowledgements

We thank Barry Levin for helpful conversations and guidance about the utilization and maintenance of the DIO (OP) and DR (OR) lines. We thank Julia Deshazor, Kimberly Gates and the UM Rodent Breeding Core for their assistance maintaining the breeding colony and generating many of the rats used in these studies. We thank Zufan Johnson, Juliette Manzur, and Edison Zheng for help with analysis of electrophysiology data, and Emaa Elrayah for help with behavioral studies. We thank Dr. Joseph J. Ziminski for his technical assistance with Easy Electrophysiology, including adapting program features to enhance analysis. This work was supported in part by NIH NIDDK R01 DK130246 to CRF and NIH NIDDK F32DK1413571 to LMR.

## Author Contributions

All authors contributed to writing of the manuscript, the conceptualization of the studies, interpretation, and aspects of data analysis and presentation; JSC conducted the majority of behavioral and whole-body physiological studies; MOD, RJC and LMR conducted electrophysiological experiments and analyzed related data.

## Figure Captions

## Notes

### Competing Interest Statement

The authors have declared no competing interest.

